# Habitat Type and Seasonal Variation as Drivers of Mosquito Proliferation in Urban Aquatic Microhabitats

**DOI:** 10.64898/2026.07.15.738763

**Authors:** S. Garrison Carruth, Jenna C. Michael, Chalmers Vasquez, John-Paul Mutebi, Maria Litvinova, André B. B. Wilke

**Affiliations:** Department of Epidemiology and Biostatistics, Indiana University School of Public Health, Bloomington, Indiana, USA; Miami-Dade County Mosquito Control Division, Miami, Florida, USA

**Keywords:** Urbanization, Mosquitoes, Vector-Borne Disease, Vector Control, Population Dynamics

## Abstract

*Aedes aegypti* is the primary vector species for dengue, Zika, and chikungunya viruses, and is adapted to thrive in urban environments. Their proximity to human populations and the adverse health outcomes associated with arboviral infections make mosquito control essential for reducing the risk of outbreaks and disease spread. Source reduction and larvicide applications are central components of mosquito control because they reduce adult mosquito emergence by targeting immature life stages. Therefore, our objective was to identify urban aquatic habitats used by *Ae. aegypti* and assess temporal variation in their contribution to immature mosquito production. To distinguish habitat use from habitat productivity, we considered larval presence as an indicator of oviposition or early-stage survival and pupal presence as evidence that a habitat supported development through most of the immature life cycle. Between July 2018 and October 2019, 2,482 inspections in Miami-Dade County, Florida, identified 2,756 aquatic habitats from which 19,466 *Ae. aegypti* larvae and 3,648 pupae were collected. Unique temporal trends were observed for both larval and pupal presence and abundance at county and habitat resolutions. County-level trends showed the highest larval and pupal production during Summer and the lowest pupal production during Winter. However, temporal patterns differed across habitats, supporting a context-dependent interpretation of habitat conduciveness, in which the same habitat type may support different levels of immature mosquito production depending on local environmental conditions. Our results show that habitats with frequent larval occurrence did not consistently have high pupal presence, indicating that larval presence does not necessarily reflect habitat productivity. Control strategies that prioritize habitats repeatedly associated with pupal production by season may improve year-round mosquito control, outbreak preparedness, and resource allocation.

## Background

*Aedes aegypti* (Diptera: Culicidae) is the primary vector species for dengue, Zika, and chikungunya viruses (Powell 2018). Dengue is the most widespread mosquito-borne arboviral disease, with an estimated 390 million infections annually worldwide (Bhatt et al. 2013). Moreover, current estimates indicate that dengue transmission could expand beyond its current endemic range (Iwamura et al. 2020; Gwee et al. 2021). Zika and chikungunya have also emerged as major public health threats, with large outbreaks in the Americas between 2014 and 2017 (Bartholomeeusen et al. 2023; Rabe et al. 2025). Preventing Zika outbreaks is especially important as it can cause microcephaly in infants whose mothers are infected during pregnancy (Antoniou et al. 2020). Beyond their substantial morbidity and mortality, these infections impose considerable economic costs, underscoring the need for effective control of *Ae. aegypti* populations (Roiz et al. 2024).

Controlling *Ae. aegypti* populations in urban areas is a challenging task that requires the coordination and implementation of multiple steps in a logical framework, often guided by Integrated Mosquito Management (IMM) (World Health Organization 2012; Lizzi et al. 2014). One of the most important and effective IMM components to control *Ae. aegypti* populations are source reduction (i.e., larvae and pupae reduction through breeding habitat elimination) and the use of larvicides (American Mosquito Control Association 2021). However, the availability of artificial and natural aquatic habitats is spatially heterogeneous across urban landscapes. Consequently, despite the ecological plasticity of *Ae. aegypti*, spatial and temporal variation in aquatic habitat availability, and colonization success drive mosquito proliferation (Powell and Tabachnick 2013).

The identification of diverse and widespread aquatic habitats that can be used as breeding sites for *Ae. aegypti* in urban environments is instrumental in guiding mosquito control strategies and effectively improving outbreak preparedness and response (Fouet and Kamdem 2019). Therefore, our objective was to identify urban aquatic habitats used by *Ae. aegypti* and assess temporal variation in their contribution to immature mosquito production. To distinguish habitat use from habitat productivity, we considered larval presence as an indicator of oviposition or early-stage survival and pupal presence as evidence that a habitat supported development through most of the immature life cycle.

## Methods

### Study Site

This study was carried out in Miami-Dade County, Florida. *Aedes aegypti* larvae and pupae were collected across 2,482 inspections from July 2018 to October 2019. Inspections were initiated by 3-1-1 complaint calls directed to the Miami-Dade Mosquito Control Division. County inspectors were dispatched to survey the area and collect immature mosquitoes in aquatic habitats within a 50-meter radius of the complaint location (Wilke et al. 2019a). These 3-1-1 calls are a normal occurrence for counties in the State of Florida and have been used previously to analyze mosquito production and control activities (Stephen T. Holmes 2007; Wilke et al. 2019a).

### Specimen Collection

Specimens were collected using manual plastic pumps and entomological dippers, depending on habitat type. Larvae and pupae were stored in plastic bags filled with 100ml of water from the aquatic habitat where they were found and transported to the Miami-Dade County Mosquito Control Laboratory for morphological identification using taxonomic keys (Darsie and Morris 2000). Larvae were identified shortly after collection, while pupae were identified after emerging as adults (Wilke et al. 2019a).

### Consolidation of habitats

Habitats were organized by: (i) mosquito presence or absence – defined as either containing immature mosquitoes (breeding site) or not (aquatic habitat only); and (ii) aquatic habitat type – defined by the physical characteristics of the reservoir collecting water allowing mosquito development. After consolidation, we selected a subset of 8 habitats that accounted for 83.3% of all *Ae. aegypti* collected for subsequent analyses. Aquatic habitats were organized as:

1. Ornamental Bromeliads – Bromeliad plants that contain phytotelmata, which provide aquatic habitats for various insects, including mosquitoes (Brown 2001).
2. Buckets – Receptacles such as barrels, buckets, or their lids/covers, primarily used in transporting or moving materials between locations.
3. Containers – Receptacles intended to hold items other than plants, including bowls, cups, jars, and boxes.
4. Drains – Drainage structures designed to remove water, including storm drains, air-conditioner drains, gutters, and runoff channels.
5. Flowerpots – Receptacles primarily intended to hold plants, as well as flowerpot accessories. This category includes dishes or plates placed under flowerpots, planters, pots, and vases.
6. Fountains – Water fixtures such as fountains and bird baths, whether functional or inactive at the time of inspection.
7. Tires – Any tire identified at a shop, household, disposal site, or discarded.
8. Trash – Garbage receptacles or litter, including garbage cans, dumpsters, trash bags, and discarded objects such as appliances.

### Population dynamics

Seasons were defined using the calendar year and northern hemisphere meteorological definitions. Summer includes June, July, and August. Fall includes September, October, and November. Winter includes December, January, and February. Spring included March, April, and May (National Oceanic and Atmospheric Administration 2024).

To assess temporal trends in immature *Ae. aegypti* populations, we calculated aquatic habitat positivity and the average number of mosquitoes collected per inspection at the county level, as well as by habitat type for each month of the collection period. For site positivity, four categories were considered:

1. Larvae-positive inspections: Proportion of all inspections in which larvae were collected in at least one aquatic habitat.
2. Larvae-positive habitats: Proportion of all aquatic habitats in which larvae were present.
3. Pupae-positive inspections: Proportion of all inspections in which pupae were collected in at least one aquatic habitat.
4. Pupae-positive habitats: Proportion of all aquatic habitats in which pupae were present.

Aquatic habitat distribution is known to be heterogeneous in urban areas (Yitbarek et al. 2023), leading to the identification of multiple habitats per inspection. Therefore, larvae and pupae positivity proportions were calculated at the county level and for each habitat by month to assess temporal trends in habitat availability, utilization, and conduciveness for immature *Ae. aegypti* proliferation.

To compare the number of immature mosquitoes over time, we calculated the average number of larvae and pupae collected per inspection for each month using the number of larvae and pupae collected as the numerator and the number of inspections per month as the denominator. This approach controlled for unequal inspection effort across the study period and enabled comparisons of immatures collected per inspection within and between habitat types over time.

### Statistical Analyses

To assess differences in observed trends in larval and pupal inspection positivity and abundance, data were grouped by season and habitat, then analyzed using four negative binomial regression models. Two negative binomial models, one for immatures collected per inspection and another for inspection positivity, were fit for larvae and for pupae. Each model had habitat and season as independent variables. To account for differences in sampling effort, the number of inspections conducted in each season was used for all models as an offset (Table 1).

**Table 1.** Statistical models used to evaluate seasonal and habitat-related variation in immature mosquito abundance and occurrence.

| **Model** | **Response variable** | **Fixed effects** | **Offset** |
| --- | --- | --- | --- |
| Larval abundance | Number of larvae collected | Season, habitat | log(seasonal inspection total) |
| Larval positivity | Number of larva-positive inspections | Season, habitat | log(seasonal inspection total) |
| Pupal abundance | Number of pupae collected | Season, habitat | log(seasonal inspection total) |
| Pupal positivity | Number of pupa-positive inspections | Season, habitat | log(seasonal inspection total) |

Negative binomial regression models were selected due to overdispersion in both the larvae and pupae data, making it a more appropriate method than implementing a model with a Poisson distribution (Zhang et al. 2025).

In each model, Winter was chosen as the reference category among seasons, and Tires were chosen for habitats. Winter was considered a baseline due to lower mosquito activity and, therefore, was selected because it had the lowest relative abundance of *Ae. aegypti* larvae and pupae. Tires was selected due to their status as an important breeding habitat (Hawley et al. 1987; Reiter 1998; Yee et al. 2010; Wilke et al. 2019b). Outliers were identified as any observation beyond the third quartile plus 1.5 times the Interquartile Range (IQR), consistent with Tukey’s method (Tukey 1977). However, these outliers were not removed from the analysis as they were considered biologically relevant for mosquito proliferation in the study area. Bee-swarm Boxplots were created to visually assess the distributional shape of immature mosquitoes collected by both season and habitat, and data were log-transformed to improve visualization. All analyses were done using R version 4.5.1. Model diagnostics were completed using the DHARMa package. Final models were conducted using the MASS package, and all plots were made using the GGPlot2 package.

## Results

A total of 2,482 inspections were conducted, resulting in 2,756 individual habitats identified, and a total of 19,466 *Ae. aegypti* larvae and 3,648 pupae collected. On average, 7.84 *Ae. aegypti* larvae and 1.47 pupae were collected per inspection. Overall, larvae were detected in 63% of inspections, whereas pupae were detected in 30%. Habitat positivity had a similar pattern, with larvae detected in 57% of all habitats inspected and in 27% of all habitats inspected for pupae. Variations in the habitat positivity and specimen abundance of immature *Ae. aegypti* was observed by season and habitat in both the larvae and pupae populations (Table 2).

**Table 2.**
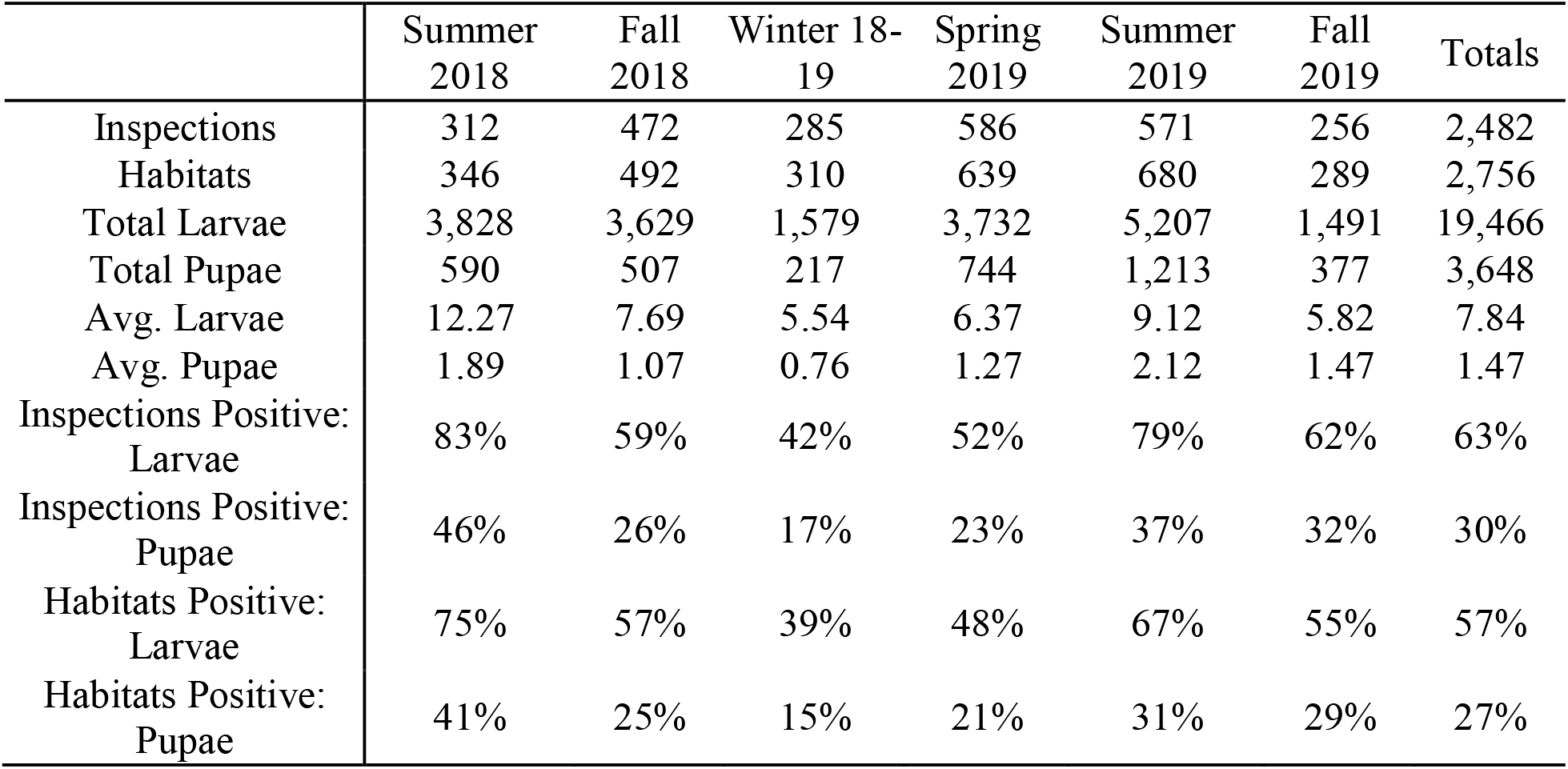
Seasonal summary of inspections and mosquito collections.

### Larvae inspection positivity

Inspection- and habitat-level positivity varied by season at the county level, peaking in June through August and reaching its lowest levels in December and January, with other months serving as transitional periods. However, February 2019 showed higher-than-average positivity of 57% among inspections and 52% among habitats (i.e., higher proportion of immature *Ae. aegypti* found in aquatic habitats inspected) compared to December 2018 and January 2019 (Figure 1a). This is 15% and 13% higher than the season average, respectively.

**Figure 1.**
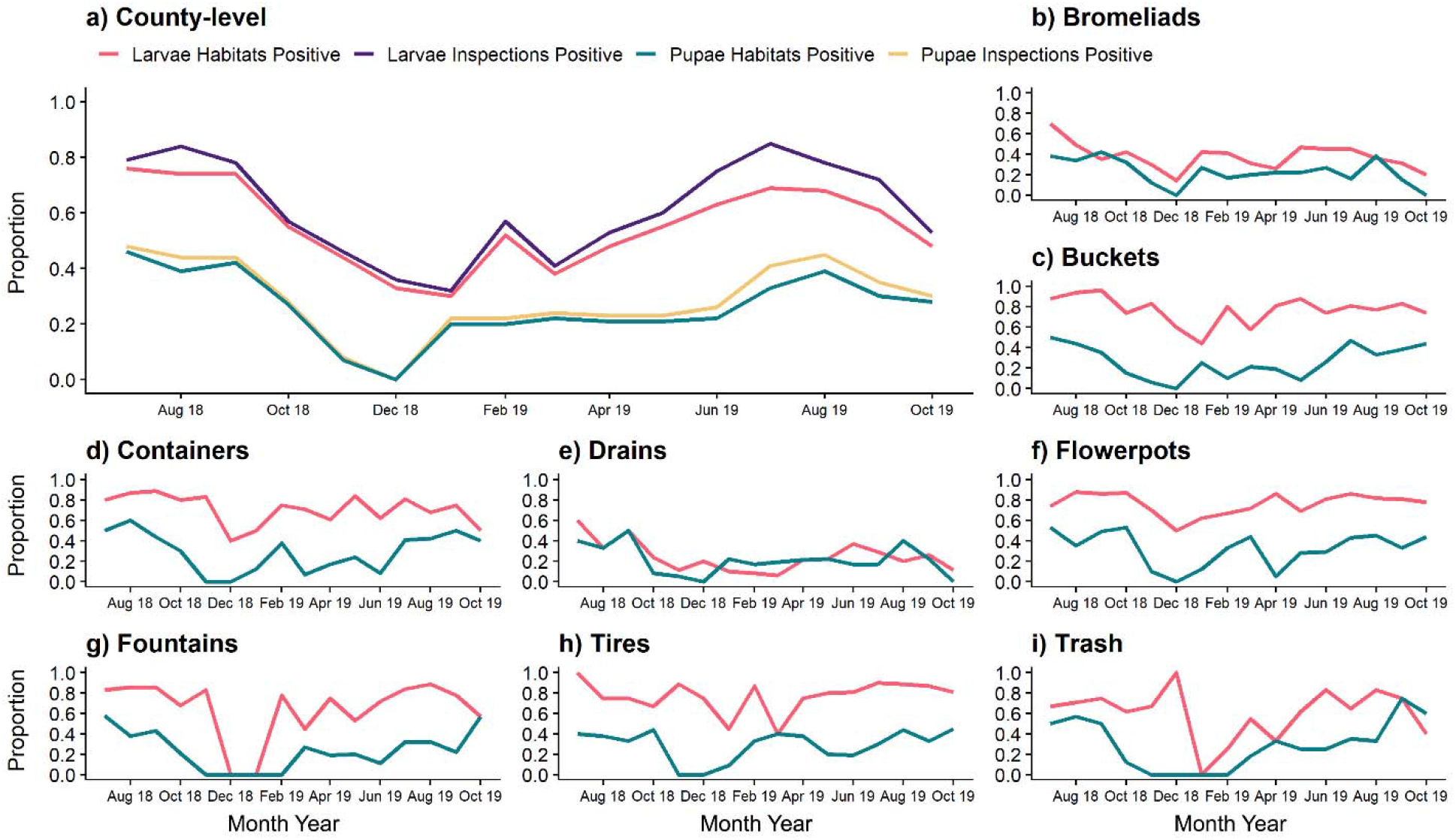
Proportion of inspections and habitats positive for immature *Aedes aegypti* between July 2018 and Oct 2019. (a) County-level proportion of larvae and pupae found by inspection (larvae = purple, pupae = gold) and habitat (larvae = red, pupae = green). (b-i) Proportion of larvae (red) and pupae (green) by habitat type.

These trends were not uniform when assessing within-habitat variation. Buckets, Containers, Flowerpots, and Tires had consistently high positivity and low monthly variation throughout the study period. Other habitats exhibited either consistent low monthly positivity, like Ornamental Bromeliads, or higher degrees of monthly variation like Drains, Fountains, and Trash. Among all aquatic habitats, Tires exhibited the highest single-month positivity, with 100% of Tires inspected containing larvae in July 2018, followed by Buckets, with 96% testing positive for larval *Ae. aegypti* in September 2018. Flowerpots were the most consistent aquatic habitat, with the highest minimum monthly positivity of 50%, followed by Buckets at 44%.

Most aquatic habitats reached their lowest positivity between December and February. Fountains, an otherwise consistently positive habitat, expeienced substantial declines with no larvae collected in December 2018 and January 2019. Drains remained consistently low across the study period, with monthly positivity ranging from 60% in July 2018 to 6% in March 2019. Ornamental Bromeliads also showed relatively low positivity compared with other habitat types, ranging from 70% to 14%. While positivity declined for most habitats in December 2018, Trash reached its peak during this period, followed by its lowest point in January 2019 (Figure 1b–i).

### Pupae inspection positivity

Similar to the pattern observed in larvae, inspection- and habitat-level positivity for pupae varied at the county level by season. Peak positivity was observed between June and August for both 2018 and 2019, lowest positivity between December and February, and transitional periods were observed during other months. No monthly outliers were observed relative to the seasonal average from January to June 2019, and positivity showed little variation, remaining at approximately 24% throughout this period (Figure 1a).

A high level of heterogeneity in pupae positivity was observed by aquatic habitat type. No specific habitat type yielded consistent positivity. However, all habitat types had at least one month with 0% positivity and at least one month exceeding 40% positivity. Most habitats had a lower inspection positivity of pupal *Ae. aegypti* between December and February. Containers, Flowerpots, and Tires had multiple peaks throughout the study period, while other habitats followed a unimodal pattern, peaking sometime between June and August. Trash had the highest single-month pupal positivity at 75% in September 2019, followed by Containers at 60% in August 2018 (Figure 1b–i).

### Larvae Averages per Inspection

Larval averages per inspection had a less defined temporal pattern than larval inspection and habitat positivity. The county-level average per inspection was unimodal, peaking in summer during July 2018 (16.16 larvae per inspection) and July 2019 (10.91 larvae per inspection). However, a distinct decrease was detected in multiple months within different seasons, yielding a less defined pattern with higher levels of variation (Figure 2a). When trends were examined by habitat type, more distinct seasonal patterns were observed than at the county level.

**Figure 2.**
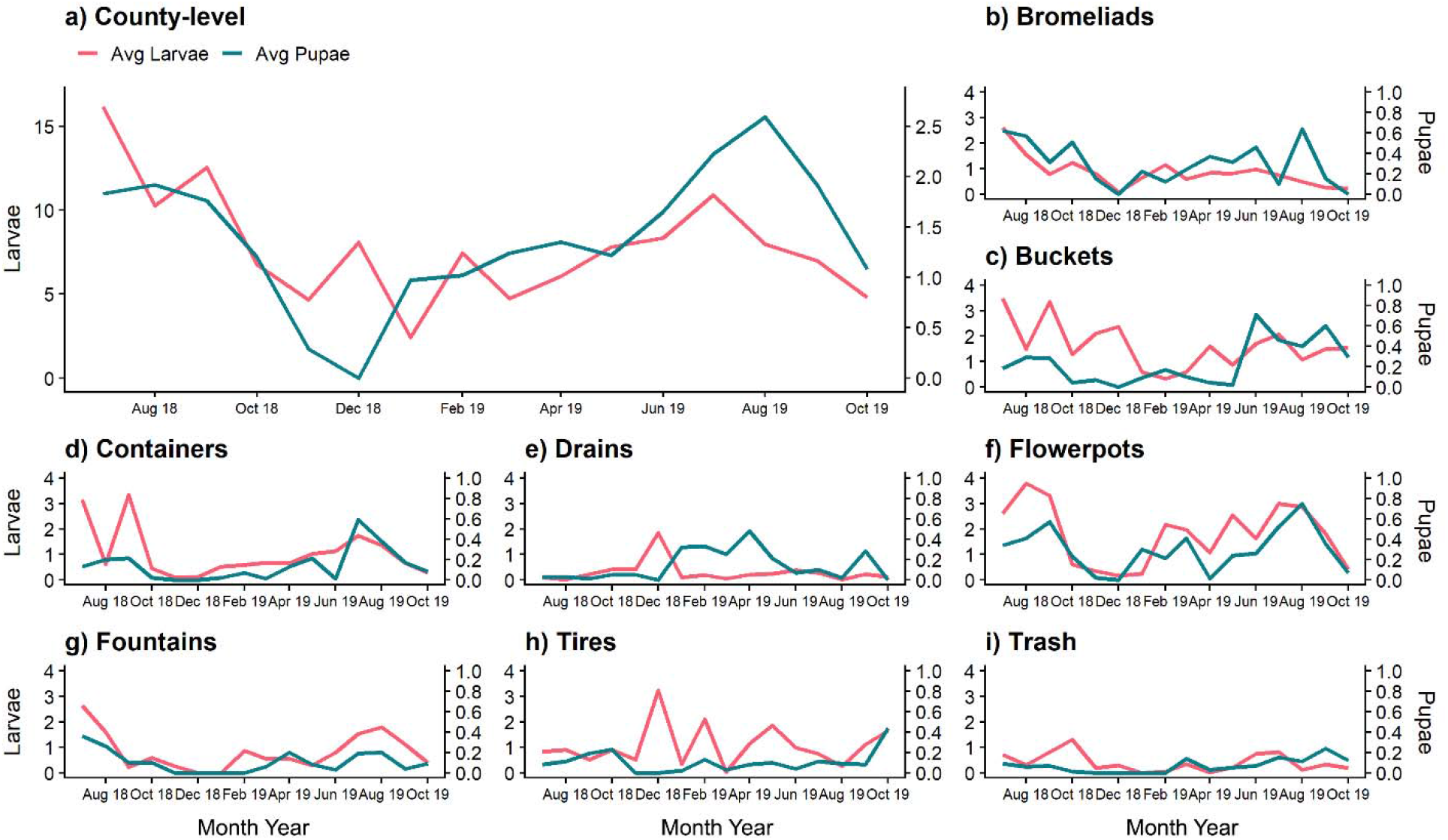
Monthly average of immature mosquitoes collected between July 2018 and Oct 2019. (a) County-level averages of larvae (red) and pupae (green) collected. (b-i) Within habitat averages.

Among habitats, Ornamental Bromeliads showed a consistent per inspection average of larvae collected throughout the study period, peaking in July and August 2018. Containers and Fountains yielded a similar pattern, with high numbers collected per inspection between June and August each year. Buckets and Flowerpots had the most distinct population dynamics, although their periods of lowest averages occurred at different times. Buckets reached their low point in February 2019, whereas Flowerpots reached their low point in December 2018 and January 2019. Drains and Tires had a unique pattern compared to other habitat types, peaking in December 2018, when most aquatic habitats had their lowest per inspection collection levels. Trash peaked in October 2018 and maintained relatively low production per inspection for the rest of the study period compared to other habitat types (Figure 2b-i).

### Pupae Averages per Inspection

Pupal per inspection averages had a clear seasonality over the study period. At the county level, a unimodal trend was observed, peaking in August 2018 (1.92 pupae per inspection) and in August 2019 (2.59 pupae per inspection). Moreover, a well-defined decrease in the number of pupae per inspection was observed in December 2018, during which no pupae were collected (Figure 2a).

Interestingly, this pattern was not uniformly present within habitat type. Only Ornamental Bromeliads largely followed the county-level pattern. Buckets and Containers maintained consistent per inspection averages until increases were observed in 2019 from June to August. Fountains, Tires, and Trash yielded consistently low levels throughout the study period. Drains were also unimodal, showing a period of highest average per inspection between January 2018 and April 2019, opposite to the trend observed at the county level. Flowerpots showed the greatest variation with three distinct peaks in per inspection average in September 2018, March 2019, and August 2019. Flowerpots also had the highest overall average, with 0.75 pupae collected per inspection (Figure 2b-i).

### Statistical Analysis

Inspection positivity varied seasonally and between habitat types for both *Ae. aegypti* larvae and pupae. For both development stages, Summer and Fall exhibited significantly higher incidence rates of positive inspections relative to Winter. By Habitat, incidence rates of positive inspections were not uniform between development stages. For larvae, all habitats except Containers and Fountains differed significantly from Tires, with Ornamental Bromeliads, Buckets, and Flowerpots having higher incidence rates, while Drains and Trash had a significantly lower incidence. For pupae, only two Habitats differed significantly from Tires, with Ornamental Bromeliads having a higher incidence and Trash showing a lower incidence (Table 3).

**Table 3.**
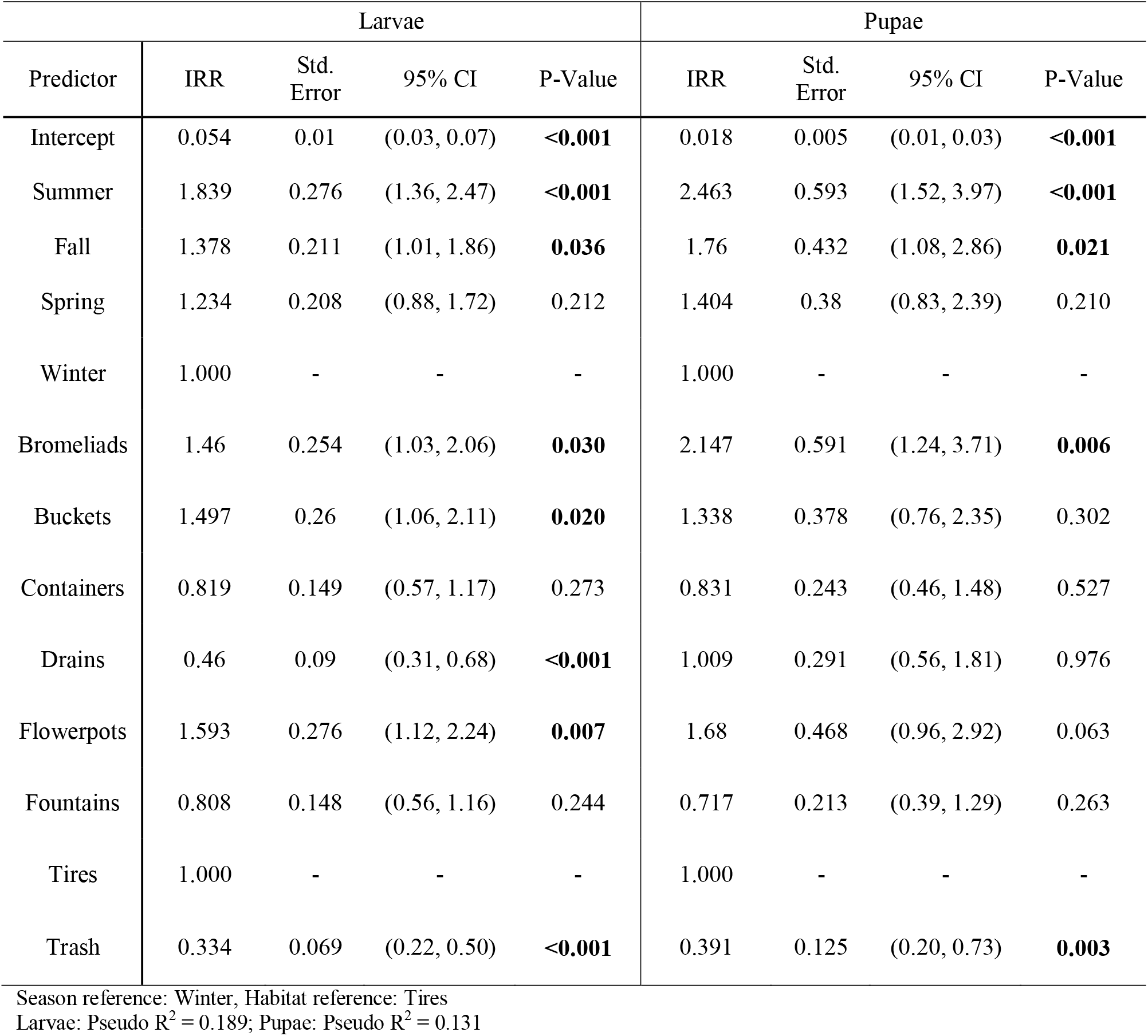
Incidence rate ratios for larval and pupal habitat positivity.

There were also seasonal and between-habitat differences in the number of both *Ae. aegypti* larvae and pupae collected. While the monthly variation of larval inspection averages showed unique temporal trends by habitat, when assessing overall differences to Tires, larval counts showed uniformity. Flowerpots and Buckets were the only habitats with increased incidence rates relative to Tires; however, the differences were not significant. Significant differences were only observed within Drains and Trash, which had 75.9% and 67.1% lower incidence rates of larvae than Tires, respectively. By season, only Summer differed statistically from Winter, with a 68.7% higher incidence of larvae (Table 4).

**Table 4.** Seasonal and habitat-related differences in larval and pupal specimen count.

| Predictor | Larvae |  |  |  | Pupae |  |  |  |
| --- | --- | --- | --- | --- | --- | --- | --- | --- |
|  | IRR | Std. Error | 95% CI | P-Value | IRR | Std. Error | 95% CI | P-Value |
| Intercept | 0.917 | 0.238 | (0.57, 1.53) | 0.739 | 0.058 | 0.021 | (0.03, 0.12) | **<0.001** |
| Summer | 1.687 | 0.365 | (1.06, 2.62) | **0.016** | 3.067 | 0.912 | (1.62, 5.65) | **<0.001** |
| Fall | 1.168 | 0.253 | (0.74, 1.79) | 0.474 | 2.080 | 0.621 | (1.10, 3.82) | **0.014** |
| Spring | 0.965 | 0.241 | (0.58, 1.59) | 0.887 | 2.027 | 0.691 | (1.02, 4.02) | **0.038** |
| Winter | 1.000 | - | - | - | 1.000 | - | - | - |
| Bromeliads | 0.74 | 0.212 | (0.41, 1.31) | 0.294 | 2.341 | 0.907 | (1.07, 5.11) | **0.028** |
| Buckets | 1.331 | 0.381 | (0.75, 2.35) | 0.319 | 1.850 | 0.718 | (0.85, 4.02) | 0.113 |
| Containers | 0.789 | 0.226 | (0.44, 1.39) | 0.408 | 1.010 | 0.394 | (0.46, 2.22) | 0.979 |
| Drains | 0.241 | 0.07 | (0.13, 0.42) | **<0.001** | 1.469 | 0.571 | (0.65, 3.30) | 0.322 |
| Flowerpots | 1.513 | 0.433 | (0.85, 2.68) | 0.148 | 2.340 | 0.907 | (1.07, 5.10) | **0.028** |
| Fountains | 0.672 | 0.193 | (0.37, 1.19) | 0.166 | 0.782 | 0.306 | (0.35, 1.73) | 0.530 |
| Tires | 1.000 | - | - | - | 1.000 | - | - | - |
| Trash | 0.329 | 0.095 | (0.18, 0.58) | **<0.001** | 0.549 | 0.217 | (0.25, 1.20) | 0.128 |

Season reference: Winter, Habitat reference: Tires
Larvae: Pseudo $R^2 = 0.082$ ; Pupae: Pseudo $R^2 = 0.070$

Pupal specimen counts by season showed more distinct seasonal variation than larvae. All Seasons were significantly different when compared to Winter. Summer had a 207% higher incidence rate of pupae relative to Winter, with Spring and Fall exhibiting more modest incidence rate increases of 103% and 108%, respectively. This highlights a clear pattern with Summer producing the highest specimen counts of *Ae. aegypti* pupae, while Fall and Spring serve as transitional periods before a yearly low during Winter. Among habitats, both Ornamental Bromeliads and Flowerpots showed significantly higher incidences of pupae relative to Tires (Table 4).

Specimen counts for larvae have shown a seasonal pattern with higher medians for the summer months and a lower median for the winter months. Winter also had the narrowest upper-quartile range. Based on Tukey’s fences, 30.8% of positive inspections were classified as outliers; however, larvae abundance was not predominantly driven by outliers. Specimen counts for pupae were primarily driven by outlier inspections for all Seasons except Summer 2018 (21.83% of positive inspections), with all having a median collection count of zero. Summers had the largest upper quartile ranges and Tukey’s fence limits, followed by Fall 2018 and 2019, while Winter and Spring’s counts were entirely driven by outliers (Figure 3b, Table 5).

**Figure 3.**
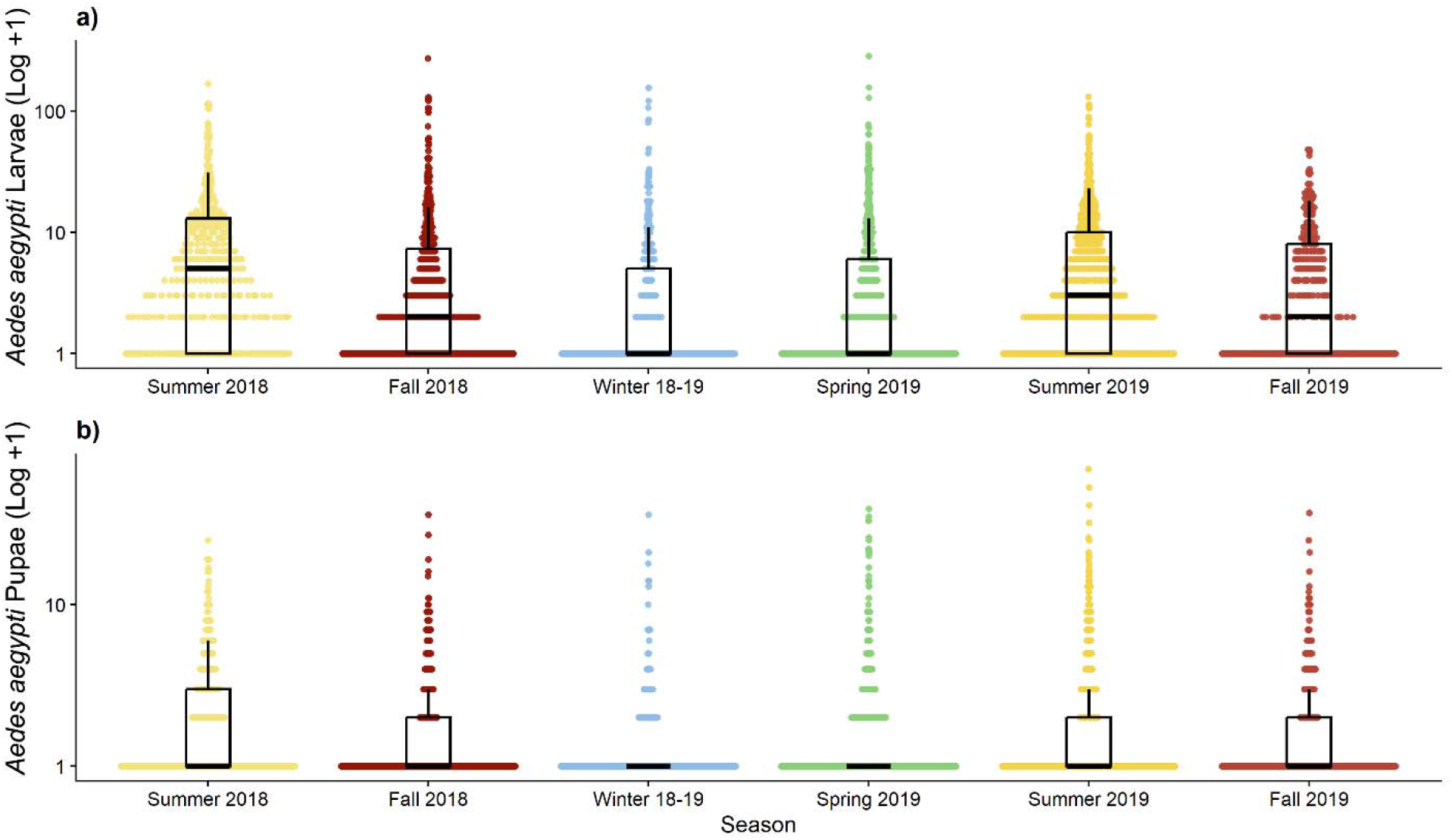
Immature mosquito collections of *Aedes aegypti* per season from Summer 2018 to Fall 2019. Log-scaled distribution of larvae (a) and pupae (b) collected per habitat inspected within each season.

**Table 5.**
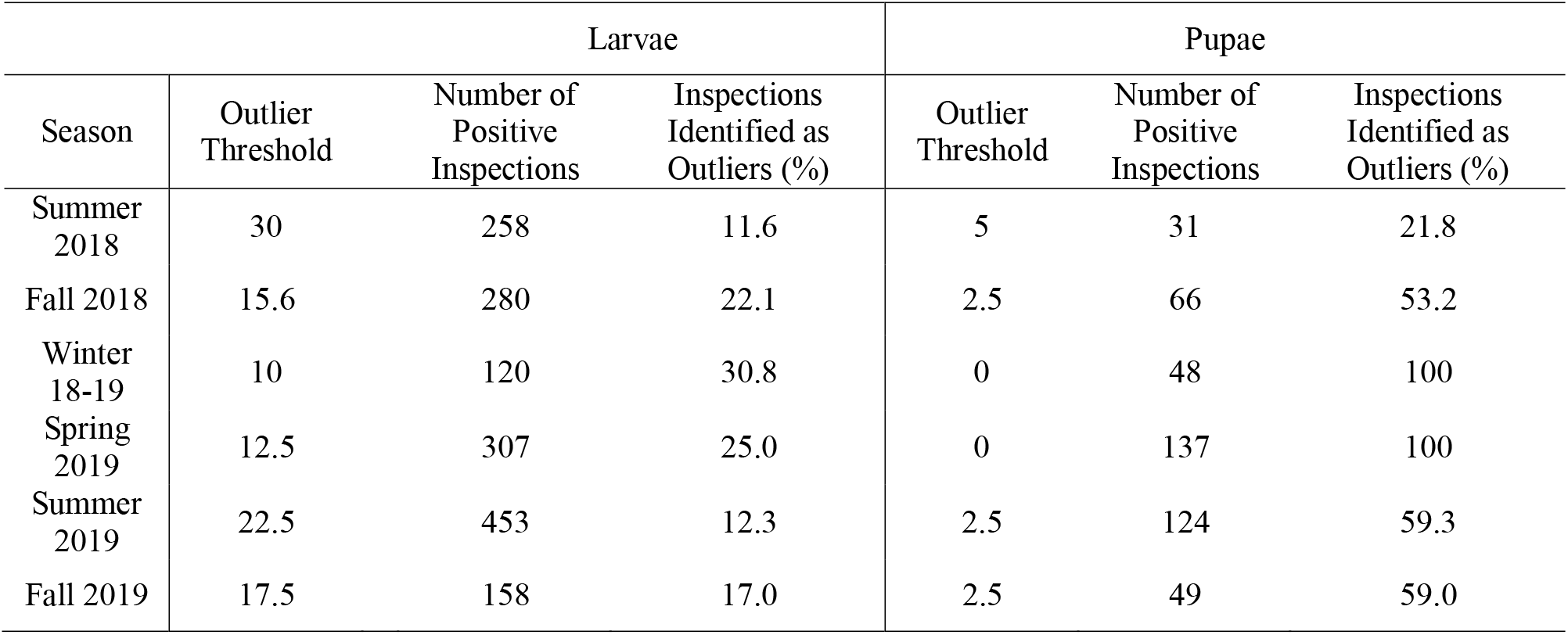
Assessment of larval and pupal counts accounted for by outlier inspection counts by season.

Although 2,791 larvae were collected from Ornamental Bromeliads and Drains, their median larval specimen counts were zero, with counts primarily driven by outlier inspections at 41.2% and 100% of positive inspections, respectively. In contrast, the median specimen count for the other habitat types ranged from two larvae per inspection for Trash to five per inspection in both Buckets and Flowerpots, with larvae counts primarily driven by non-outlier inspections according to Tukey’s outlier detection method. (Figure 4a, Table 6). Outlier inspections within habitats mostly drove pupae production. Outliers did not drive pupae production in Flowerpots (27% of positive pupae inspections) and Tires (44.4% of positive pupae inspections), whereas in other habitat types, most inspections containing pupae fell within the outlier range. Pupae counts of Ornamental Bromeliads and Drains were exclusively driven by outliers (Figure 4b, Table 6).

**Figure 4.**
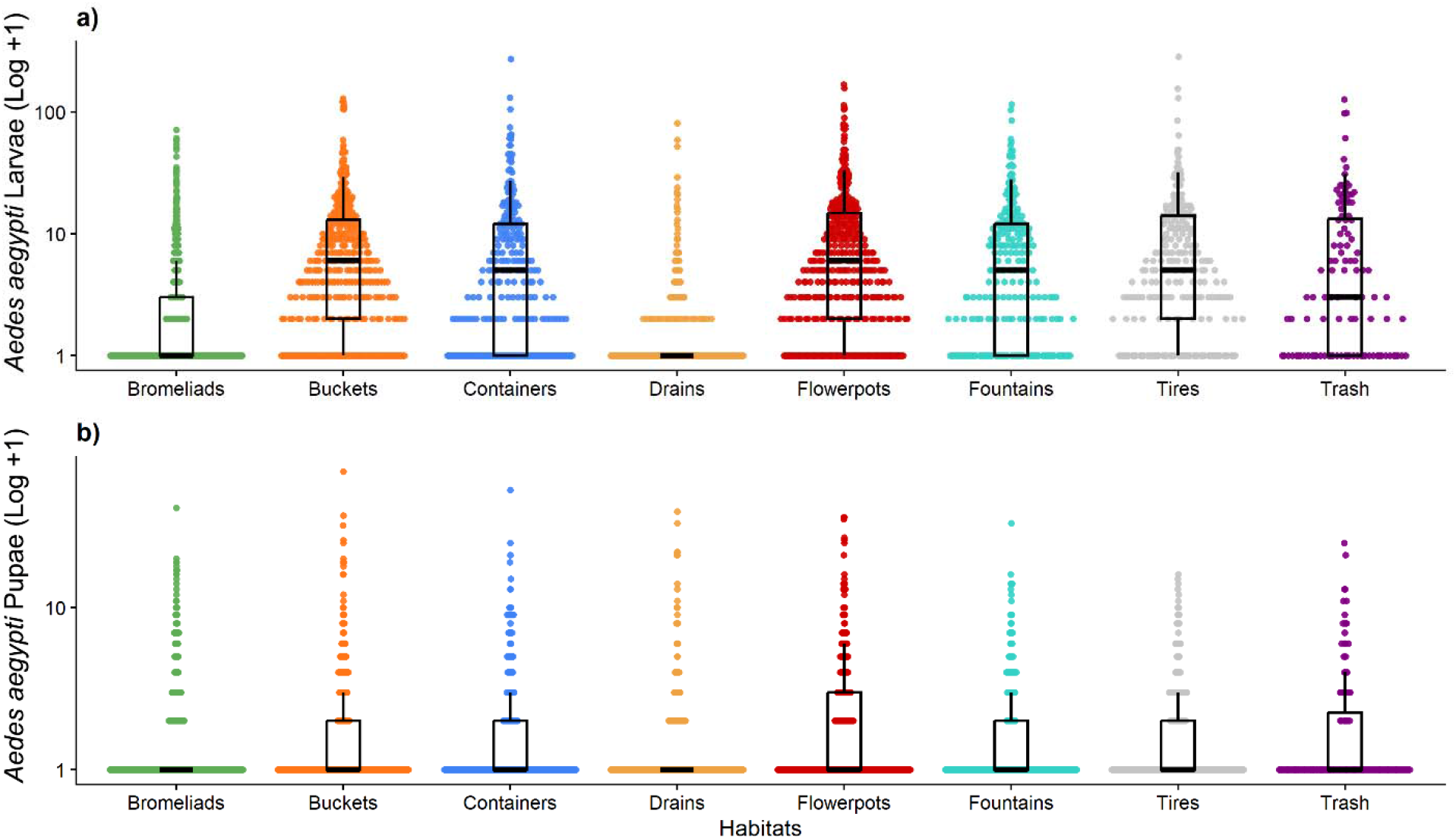
Immature mosquito collections of *Aedes Aegypti* per habitat from Summer 2018 to Fall 2019. Log-scaled distribution of larvae (a) and pupae (b) collected per inspection within each habitat.

**Table 6.** Assessment of larval and pupal specimen counts accounted for by outlier inspection counts by habitat.

| Habitat | Larvae |  |  | Pupae |  |  |
| --- | --- | --- | --- | --- | --- | --- |
|  | Outlier Threshold | Number of Positive Inspections | Inspections Identified as Outliers (%) | Outlier Threshold | Number of Positive Inspections | Inspections Identified as Outliers (%) |
| Bromeliads | 5 | 288 | 41.2 | >0 | 179 | 100 |
| Buckets | 28.5 | 292 | 10.9 | 2.5 | 108 | 61.1 |
| Containers | 27.5 | 169 | 12.4 | 2.5 | 70 | 60 |
| Drains | >0 | 87 | 100 | >0 | 72 | 100 |
| Flowerpots | 32.8 | 330 | 9.0 | 5 | 148 | 27.0 |
| Fountains | 27.5 | 165 | 10.9 | 2.5 | 61 | 62.3 |
| Tires | 31 | 176 | 7.3 | 2.5 | 72 | 44.4 |
| Trash | 30.6 | 69 | 8.7 | 3.1 | 33 | 51.5 |

## Discussion

Urban landscapes contain thousands of small, discontinuous aquatic habitats embedded within highly anthropized environments. However, a major limitation of many habitat-productivity frameworks is the implicit assumption that specific habitat types have stable productivity values. Our results show clear habitat-specific temporal variation in both the number of aquatic habitats supporting immature *Ae. aegypti*, and the number of immature *Ae. aegypti* specimens per habitat.

The seasonal pattern observed at the county level suggests that climate and seasonality define a broad temporal envelope for immature *Ae. aegypti* occurrence. Larval and pupal positivity were highest during summer and lowest during winter, with spring and fall representing intermediate periods. This pattern was also reflected in the statistical models, in which summer and fall had higher incidence rates of positive inspections than winter for both larvae and pupae. Pupal abundance showed a stronger seasonal pattern than larval abundance, with summer, fall, and spring differing from winter. These findings are consistent with previous studies showing that temperature and rainfall influence immature mosquito development (Troyo et al. 2008; Egid et al. 2022; Prasad et al. 2024; Soria et al. 2025).

However, the habitat-specific patterns observed in this study indicate that temperature and rainfall did not drive realized immature *Ae. aegypti* production alone. Larval and pupal presence and abundance patterns differed across habitats, indicating that habitats where larvae occurred were not always the same habitats where high pupal production was observed. For larvae, Buckets and Flowerpots showed higher counts across the study period; Bromeliads, Containers, and Tires showed intermittent periods of higher larval abundance; and Drains, Fountains, and Trash had overall lower larval counts. For pupae, Bromeliads and Flowerpots accounted for higher pupal counts, whereas Buckets, Containers, and Drains showed intermittent pupal production, and Fountains, Tires, and Trash yielded lower overall pupal counts. These immature *Ae. aegypti* developmental stage-specific patterns support a potential context-dependent interpretation of habitat conduciveness, in which realized immature mosquito production varies by habitat type, season, and local environmental conditions. Larval presence indicates oviposition or early-stage survival, whereas pupal abundance provides stronger evidence that an aquatic habitat supported development through most of the immature life cycle (Midega et al. 2006; Chadee et al. 2009).

High-count aquatic habitats contributed substantially to the observed immature *Ae. aegypti* counts, particularly for pupae. Larval counts were more broadly distributed across aquatic habitats, whereas pupal counts were concentrated in a smaller subset of habitats, frequently above the outlier threshold set by Tukey’s formula. This pattern is consistent with local environmental filtering during immature development, where many habitats support colonization or early-stage survival, but fewer support development to the pupal stage.

Habitat conduciveness also varied locally. For example, our results show that Ornamental Bromeliads were conducive to *Ae. aegypti* production in Miami-Dade, conflicting with previous studies carried out in parts of Africa, Brazil, and Sri Lanka, in which natural habitats such as Ornamental Bromeliads are less relevant to *Ae. aegypti* proliferation (Mocellin et al. 2009; Ngugi et al. 2017; Egid et al. 2022; Herath et al. 2024). This difference supports a context-dependent interpretation of habitat conduciveness, in which the same habitat type may support different levels of immature mosquito production depending on specific local environmental conditions (Barrera et al. 2011; Islam et al. 2019; Egid et al. 2022; Mwakutwaa et al. 2023; Avramov et al. 2024; Prasad et al. 2024; Zanoni et al. 2025). Differences between larval occurrence and pupal production are consistent with a developmental bottleneck, in which local environmental filters limit progression from larvae to pupae in aquatic habitats (Troyo et al. 2008; Wilke et al. 2018; Prasad et al. 2024).

These results are also relevant for mosquito control and arbovirus outbreak preparedness and response. Larval metrics alone are insufficient to predict pupal abundance and subsequent adult mosquito production necessary to guide and support mosquito control operations (Midega et al. 2006; Chadee et al. 2009). Because pupae provide a closer indication of adult mosquito emergence, habitat assessments that incorporate both larval and pupal metrics can better identify aquatic habitats contributing to realized mosquito production that could be prioritized in mosquito control strategies. Our findings support habitat- and season-specific control strategies that distinguish habitats with frequent larval occurrence from those that repeatedly support pupal production. Such targeted approaches may improve the allocation of mosquito control effort and reduce reliance on broad, non-specific larvicide applications (Egid et al. 2022; Soria et al. 2025).

During this study, we relied on a heterogeneous sampling effort. Therefore, sampling effort was normalized by calculating monthly immature mosquito abundance as the number of larvae or pupae collected per inspection. Additionally, to account for the heterogeneous sampling effort in the regression models, the number of inspections conducted during each season was included as an offset. Regression analysis did not consider potential confounders to the relationship between habitat positivity or abundance and seasonality, and habitat type. Moreover, interaction terms were not possible due to limitations in model stability.

Further research is needed to evaluate aquatic habitats in urban landscapes as assemblages rather than isolated units. These co-occurring habitats may create complementary developmental opportunities, with some maintaining water during dry periods, others accumulating organic matter after rainfall, and others providing protection from the elements. Such habitat assemblages could generate more consistent or higher immature mosquito production than expected from individual habitats alone, but these potential non-additive effects remain poorly understood.

## Conclusion

This study identified urban aquatic habitats used by *Ae. aegypti* and assessed temporal variation in their contribution to immature mosquito production. Our results show that habitats with frequent larval occurrence did not consistently show high pupal presence, indicating that oviposition or early-stage survival does not necessarily translate into successful development through the immature life cycle. These findings support distinguishing habitat use from habitat productivity in *Ae. aegypti* surveillance and control. Larval surveys alone may not adequately identify habitats contributing most to adult mosquito emergence, particularly when local environmental conditions affect survival from larval to pupal stages. Habitat- and season-specific control strategies that prioritize sites repeatedly associated with pupal production may improve year-round mosquito control, outbreak preparedness, and resource allocation.

## Funding

S.G.C., J.C.M., M.A., and A.B.B.W. were supported by the National Science Foundation (DMS-2526926). The funder had no role in the design of the study and collection, analysis, and interpretation of data, and in writing the manuscript.

## Data Availability

Data from Miami-Dade County analyzed in this study were obtained from a publicly supported surveillance program. Although such data are public property and may ultimately become part of the public record, access is not automatically unrestricted or immediate. Individuals seeking access to the surveillance data used in this study must obtain permission from the appropriate public health authority to ensure compliance with applicable public-records statutes, data-use policies, and privacy protections. Researchers interested in obtaining the dataset may submit a formal Public Records Request to: Miami-Dade County Mosquito Control Division, available at: https://www.miamidade.gov/global/solidwaste/mosquito/contact-mosquito-control.page.

## Notes

### Competing Interest Statement

The authors have declared no competing interest.

## References

American Mosquito Control Association (2021) Best practices for integrated mosquito management. Sacramento, CA

Antoniou E, Orovou E, Sarella A, Iliadou M, Rigas N, Palaska E, Iatrakis G, Dagla M (2020) Zika virus and the risk of developing microcephaly in infants: A systematic review. Int J Environ Res Public Health 17:11. 10.3390/ijerph17113806

Avramov M, Thaivalappil A, Ludwig A, Miner L, Cullingham CI, Waddell L, Lapen DR (2024) Relationships between water quality and mosquito presence and abundance: A systematic review and meta-analysis. J Med Entomol 61:1–33. 10.1093/jme/tjad139

Barrera R, Amador M, MacKay AJ (2011) Population dynamics of *Aedes aegypti* and dengue as influenced by weather and human behavior in San Juan, Puerto Rico. PLoS Negl Trop Dis 5:e1378. 10.1371/journal.pntd.0001378

Bartholomeeusen K, Daniel M, LaBeaud DA, Gasque P, Peeling RW, Stephenson KE, Ng LFP, Ariën KK (2023) Chikungunya fever. Nat Rev Dis Primers 9:17. 10.1038/s41572-023-00429-2

Bhatt S, Gething PW, Brady OJ, Messina JP, Farlow AW, Moyes CL, Drake JM, Brownstein JS, Hoen AG, Sankoh O, Myers MF, George DB, Jaenisch T, Wint GRW, Simmons CP, Scott TW, Farrar JJ, Hay SI (2013) The global distribution and burden of dengue. Nature 496:504–507. 10.1038/nature12060

Brown B V. (2001) Flies, gnats, and mosquitoes. Encyclopedia of Biodiversity 815–826. 10.1016/B0-12-226865-2/00123-1

Chadee DD, Huntley S, Focks DA, Chen AA (2009) *Aedes aegypti* in Jamaica, West Indies: Container productivity profiles to inform control strategies. Tropical Medicine & International Health 14:220–227. 10.1111/j.1365-3156.2008.02216.x

Darsie RF, Jr., Morris CD (2000) Keys to the adult females and fourth-instar larvae of the mosquitoes of Florida. 1st ed Vol 1 Technical Bulletin of the Florida Mosquito Control Association

Egid BR, Coulibaly M, Dadzie SK, Kamgang B, McCall PJ, Sedda L, Toe KH, Wilson AL (2022) Review of the ecology and behaviour of *Aedes aegypti* and *Aedes albopictus* in Western Africa and implications for vector control. Current Research in Parasitology & Vector-Borne Diseases 2:100074. 10.1016/j.crpvbd.2021.100074

Fouet C, Kamdem C (2019) Integrated mosquito management: Is precision control a luxury or necessity? Trends Parasitol 35:85–95. 10.1016/j.pt.2018.10.004

Gwee XWS, Chua PEY, Pang J (2021) Global dengue importation: A systematic review. BMC Infect Dis 21:1078. 10.1186/s12879-021-06740-1

Hawley WA, Reiter P, Copeland RS, Pumpuni CB, Craig GB, Jr. (1987) *Aedes albopictus* in North America: Probable introduction in used tires from Northern Asia. Science (1979) 236:1114–1116. 10.1126/science.3576225

Herath JMMK, De Silva WAPP, Weeraratne TC, Karunaratne SHPP (2024) Breeding habitat preference of the dengue vector mosquitoes *Aedes aegypti* and *Aedes albopictus* from urban, semiurban, and rural areas in Kurunegala District, Sri Lanka. J Trop Med 2024:4123543. 10.1155/2024/4123543

Islam S, Haque CE, Hossain S, Rochon K (2019) Role of container type, behavioural, and ecological factors in *Aedes* pupal production in Dhaka, Bangladesh: An application of zero-inflated negative binomial model. Acta Trop 193:50–59. 10.1016/j.actatropica.2019.02.019

Iwamura T, Guzman-Holst A, Murray KA (2020) Accelerating invasion potential of disease vector Aedes aegypti under climate change. Nat Commun 11:2130. 10.1038/s41467-020-16010-4

Lizzi KM, Qualls WA, Brown SC, Beier JC (2014) Expanding integrated vector management to promote healthy environments. Trends Parasitol 30:394–400. 10.1016/j.pt.2014.06.001

Midega JT, Nzovu J, Kahindi S, Sang RC, Mbogo C (2006) Application of the pupal/demographic-survey methodology to identify the key container habitats of *Aedes aegypti* (L.) in Malindi district, Kenya. Ann Trop Med Parasitol 100:61–72. 10.1179/136485906X105525

Mocellin MG, Simões TC, Nascimento TFS do, Teixeira MLF, Lounibos LP, Oliviera, RL de (2009) Bromeliad-inhabiting mosquitoes in an urban botanical garden of dengue endemic Rio de Janeiro–are bromeliads productive habitats for the invasive vectors *Aedes aegypti* and *Aedes albopictus*? Mem Inst Oswaldo Cruz 104:1171–1176. 10.1590/s0074-02762009000800015

Mwakutwaa AS, Ngugi HN, Ndenga BA, Ndenga BA, Krysosik A, Ngari M, Abubakar LU, Yonge S, Kitron U, LaBeaud AD, Mutuku FM (2023) Pupal productivity of larval habitats of *Aedes aegypti* in Msambweni, Kwale County, Kenya. Parasitol Res 122:801–814. 10.1007/s00436-022-07777-0

National Oceanic and Atmospheric Administration (2024) Infographic: Meteorological and astronomical seasons. https://www.noaa.gov/education/multimedia/infographic/infographic-meteorological-and-astronomical-seasons. Accessed 9 Jun 2026

Ngugi HN, Mutuku FM, Ndenga BA, Musunzaji PS, Mbakaya JO, Aswani P, Irungu LW, Mukoko D, Vuvule J, Kitron U, LeBeaud AD (2017) Characterization and productivity profiles of *Aedes aegypti* (L.) breeding habitats across rural and urban landscapes in western and coastal Kenya. Parasit Vectors 10:331. 10.1186/s13071-017-2271-9

Powell JR (2018) Mosquito-borne human viral diseases: Why *Aedes aegypti*? Am J Trop Med Hyg 98:1563–1565. 10.4269/ajtmh.17-0866

Powell JR, Tabachnick WJ (2013) History of domestication and spread of *Aedes aegypti* - a review. Mem Inst Oswaldo Cruz 108 Suppl 1:11–17. 10.1590/0074-0276130395

Prasad P, Gupta SK, Mahto KK, Kumar G, Rani A, Velan I, Arya DK, Singh H (2024) Influence of climatic factors on the life stages of *Aedes* mosquitoes and vectorial transmission: A review. J Vector Borne Dis 61:158–166. 10.4103/jvbd.jvbd_42_24

Rabe IB, Hills SL, Haussig JM, Walker AT, Dos Santos T, San Martin JL, Gutierrez G, Mendez-Rico J, Rodriguez JC, Elizondo-Lopez D, Gonzalez-Escobar G, Emmanuel C, Al Eryani SM, Kodama C, Yajima A, Kakkar M, Kato M, Wijesinghe PR, Samaraweera S, Brindle H, Tissera H, Kelley J, Lackritz E, Rojas DP (2025) A review of the recent epidemiology of zika virus infection. Am J Trop Med Hyg 112:1026–1035. 10.4269/ajtmh.24-0420

Reiter P (1998) *Aedes albopictus* and the world trade in used tires. 1988-1995: The shape of things to come? Journal of the American Mosquito Control Association 14:83–94

Roiz D, Pontifes PA, Jourdain F, Diagne C, Leroy B, Vaissière A, Tolsá-García MJ, Salles J, Simard F, Courchamp F (2024) The rising global economic costs of invasive *Aedes* mosquitoes and *Aedes*-borne diseases. Science of The Total Environment 933:173054. 10.1016/j.scitotenv.2024.173054

Soria C, Crocco LB, Grech MG, Stewart-Ibarra A, Almirón WR (2025) Habitat characteristics that favour the presence of *Aedes aegypti* (Diptera: Culicidae) in households in the city of Córdoba, a temperate area of Argentina. Parasit Vectors 18:487. 10.1186/s13071-025-07114-1

Stephen T. Holmes (2007) Building a 311 system: A case study of the Orange County, Florida, Government Service Center. Available at: https://www.ojp.gov/ncjrs/virtual-library/abstracts/building-311-system-case-study-orange-county-florida-government#0-0. Accessed 18 Apr 2026

Troyo A, Calderón-Arguedas O, Fuller DO, Solano ME, Avendaño A, Arheart KL, Chadee DD, Beier JC (2008) Seasonal profiles of *Aedes aegypti* (Diptera: Culicidae) larval habitats in an urban area of Costa Rica with a history of mosquito control. Journal of Vector Ecology 33:76–88. 10.3376/1081-1710(2008)33[76:SPOAAD]2.0.CO;2

Tukey J (1977) Exploratory data analysis. Addison-Wesley, Reading, Massachusetts

World Health Organization. Handbook for integrated vector management. World Health Organization; 2012.

Wilke ABB, Chase C, Vasquez C, Carvajal A, Medina J, Petrie WD, Beier JC (2019a) Urbanization creates diverse aquatic habitats for immature mosquitoes in urban areas. Sci Rep 9:15335. 10.1038/s41598-019-51787-5

Wilke ABB, Vasquez C, Mauriello PJ, Beier JC (2018) Ornamental bromeliads of Miami-Dade County, Florida are important breeding sites for *Aedes aegypti* (Diptera: Culicidae). Parasit Vectors 11:283. 10.1186/s13071-018-2866-9

Wilke ABB, Vasquez C, Petrie W, Beier JC (2019b) Tire shops in Miami-Dade County, Florida are important producers of vector mosquitoes. PLoS One 14:e0217177. 10.1371/journal.pone.0217177

Yee DA, Kneitel JM, Juliano SA (2010) Environmental correlates of abundances of mosquito species and stages in discarded vehicle tires. J Med Entomol 47:53–62 10.1603/033.047.0107

Yitbarek S, Chen K, Celestin M, McCary M (2023) Urban mosquito distributions are modulated by socioeconomic status and environmental traits in the USA. Ecological Applications 33:e2869. 10.1002/eap.2869

Zanoni MM, Santos LGRO, de Oliveira AG (2025) Socio-economic effects on the temporal importance of breeding site types for *Aedes aegypti* in a Tropical Epidemic City. Zoonoses Public Health 72:747–755. 10.1111/zph.70018

Zhang W, Liu Q, Ni J, Wang J, Gong Z (2025) Negative binomial regression analysis of factors influencing the number of distinct mosquito species in Zhejiang Province, China, 2023. Sci Rep 15:10433. 10.1038/s41598-025-94288-4

